# Determining the pathogenicity of MUT gene variant by mini-gene splicing assay

**DOI:** 10.1101/2021.02.19.432054

**Authors:** Yanyun Wang, Yun Sun, Tao Jiang

## Abstract

**Objective:** To perform gene mutation splicing analysis on 12 suspected pathogenic MMUT/MUT gene mutation sites to verify the pathogenicity of the mutation.

**Methods:** Wild-type and mutant minigenes were inserted into pcMINI vector, and total 5 wild-type recombinant vectors and 12 mutant recombinant vectors were constructed. The total RNA in 293T cells was extracted after the recombinant vectors were transfected into 293T cell line. Then PCR products were detected by agarose gel electrophoresis and analyzed by sequencing.

**Results:** RT-PCR and sequencing results showed that among 12 mutations, 9 mutations (c.419T>C, 469G>T, c.470T>A, c.626dupC, c.693C>G, c.976A>G, c.1009T>C, c.1777G>T and c.1874A>C) did not affect the gene splicing, and the other 3 mutations (c.454C>T, c.421G>A and c.2125-3C>G) all affected mRNA splicing.

**Conclusion:** In recent case reports of MMUT/MUT gene mutation sites, variant of uncertain significance (VUS) variation is very common. In this study, the pathogenicity of three mutation sites is confirmed by the mini-gene method.

## Introduction

In recent years, with the rapid development of basic research in the field of genetic metabolic diseases, diagnostic methods based on biochemistry, enzymology and cytogenetics have emerged successively. In the era of post-human genome project, various sequencing technologies have developed rapidly, among which the emergence of high-throughput sequencing (also known as next-generation sequencing) technology has greatly changed the diagnosis mode of IMD (inherited metabolism diseases), especially suspected inherited metabolism diseases screened by newborn screening via MS/MS (tandem-mass spectrometry).

However, whether it is disease-targeted gene panel or whole-exome sequencing, the judgment of pathogenicity in analyte site largely depends on database collection and literature reports^[1–4]^. For example, HGMD is a database specially used to collect the pathogenic sites closely related to human genetic diseases in the published literature, and each site has PMID of the references. For clinical genetic bioinformatics disease analysis, the literature of relevant sites can be found quickly via HGMD to avoid missing the information of pathogenic sites. However, HGMD is based on the credibility of the literature, and in fact part of sites may not be the actual pathogenic sites. Some variant of uncertain significance (VUS) variation from gene detection results have not been reported in literature. Once published as an article, these pathogenic sites with unclear clinical significance will become the reference for the subsequent gene detection data, especially in the third-party testing company whose research strength is open to discuss, these pathogenic sites will be indicated as “suspected pathogenic sites”. At present, there are few studies on the “pathogenicity” of suspected pathogenic sites related to genetic metabolic diseases. We hope that more studies will be used to confirm the actual pathogenicity of these suspected sites.

MMUT/MUT (methylmalonyl-CoA mutase) gene encodes the mitochondrial enzyme methylmalonyl Coenzyme A mutase^[5]^. In humans, the product of this gene is a VitB12-dependent enzyme which catalyzes the isomerization of methylmalonyl-CoA to succinyl-CoA which is highly expressed in liver and kidney tissue. Mutations in this gene may lead to isolated methylmalonic acidemia (MMA) with autosomal recessive inheritance which can be screened by detecting propionylcarnitine (C3) in dried blood spots (DBS) with MS/MS or detecting the concentration of methylmalonic acid in urine with GC/MS (gas chromatography-mass spectrometry)^[6]^.

In 2019, the Jiangsu Provincial Neonatal Screening Quality Control Center was established in Jiangsu Province. In 2019, several cases of isolated MMA were diagnosed in Jiangsu Province, and all the detected pathogenic sites were suspected to be pathogenic or not yet reported. Since these sites had potential splice sites which were predicted by software (e.g. Human Splicing Finder), therefore, in this study, the pathogenicity of these sites are verified by the minigene method.

## Methods

1. Collection of pathogenic sites: In this study, 12 suspected pathogenic sites (Figure 1) were collected which are suspected to be pathogenic or not yet reported before.
2. Minigene construction strategy: wild-type and mutant minigenes were inserted into pcMINI vectors. The vector skeleton was pcMINI provided by Bioeagle Co., Ltd. (China).

2.1 There are 7 mutation sites in Exon2 plus a wild-type, therefore, 8 plasmids need to be constructed which construction mode are “Exon1-Intron1-MSC-Intron2-Exon3”. “Exon1-Exon2-Exon3” is the default state of wild-type.
2.2 There are 2 mutation sites in Exon4 plus a wild-type, therefore, 3 plasmids need to be constructed which construction mode are “Exon3-Intron3-MSC-Intron4-Exon5”. “Exon3-Exon4-Exon5” is the default state of wild-type.
2.3 There are one mutation site in Exon9 plus a wild-type, therefore, 2 plasmids need to be constructed which construction mode are “Exon8-Intron8-MSC-Intron9-Exon10”. “Exon8-Exon9-Exon10” is the default state of wild-type.
2.4 There are one mutation site in Exon10 plus a wild-type, therefore, 2 plasmids need to be constructed which construction mode are “Exon9-Intron9-MSC-Intron10-Exon11”. “Exon9-Exon10-Exon11” is the default state of wild-type.
2.5 There are one mutation site in Intron11 plus a wild-type, therefore, 2 plasmids need to be constructed. The sequence of Intron11 was so long that the construction mode was adapted to “Exon11-Intron11-Exon12”. “Exon11A-Exon12” is the default state of wild-type.
3. Minigene transcription analysis: To detect whether there is a difference in the splicing mode of ExonA-Exon-ExonB, recombinant vectors were transiently transfected into 293T cells using liposome according to the instructions, and the samples were collected 36h later. The total RNA in cell samples were extracted according to the instructions. After concentration determination of RNA, cDNA was synthesized with the same amount of RNA. PCR amplification was carried out by using the primers on both sides of minigene, PCR products were detected by agarose gel electrophoresis, and the sequencing was performed.

**Figure 1.**
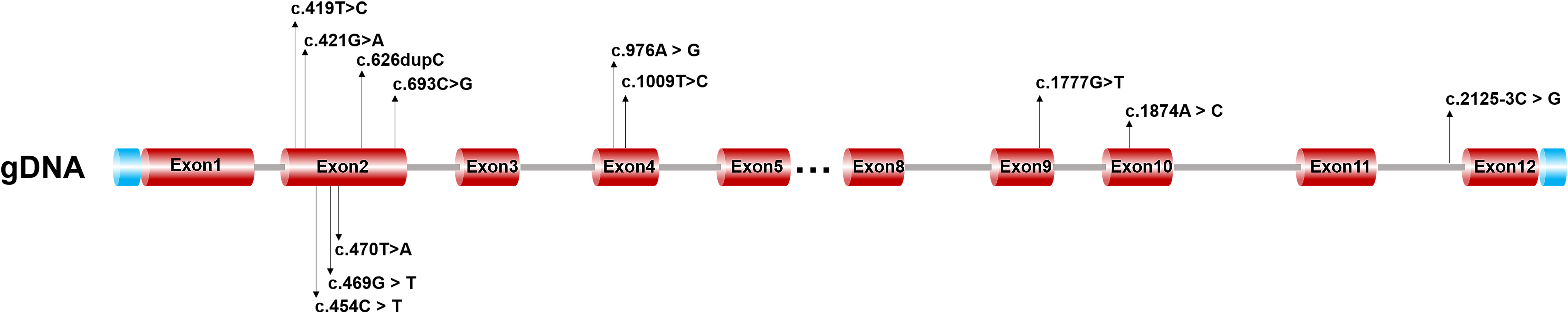
Basic information of 12 suspected pathogenic sites in MUT gene.

## Results

1. The recombined plasmids were confirmed by sequencing, and the sequencing results are shown in Figure. 2.
2. Transcription analysis results: 17 recombinant vectors (including 5 wild-type recombinant vectors and 12 mutant recombinant vectors) were transfected into 293T cells, respectively. After 36 h of transfection, a total of 17 samples were collected. RT-PCR and sequencing results were shown in Figure 3.
1. c.419T>C, 469G>T, c.470T>A, c.626dupC and c.693C>G: The wild type and the mutant type both have one band with the same size.The grayscale of normally spliced bands was scanned, and showed no difference in spliced size and splicing extent between the wild type and the mutant type. We sequenced the band of the wild type and mutant type. The sequencing results show that both wild type and mutant type were normal spliced bands and the splicing method was “Exon1-Exon2 (368bp)-Exon3”.
2. c.454C>T and c.421G>A: The wild type has one band (marked as E2-a) and the mutant type has two bands (marked as E2-a and E2-b). The sequencing results show that the small band of the wild type was normal spliced bands and the splicing method was “Exon1-Exon2 (368bp)-Exon3”. However, the large band has intron retention which retained 16bp on the right side of Intron2 as “Exon1-Exon2(368bp)- ▽ intron2(16bp)- -Exon3” . The specific splicing results are shown in Figure 3.
3. c.976A>G and c.1009T>C: The wild type and the mutant type both have one band with the same size.We sequenced the band of the wild type and mutant type. The sequencing results show that both wild type and mutant type were normal spliced bands and the splicing method was “Exon3-Exon4 (172bp)-Exon5”.
4. c.1777G>T: The wild type and the mutant type both have three bands (marked as E9-a, E9-b, E9-c) with the same size. Although there were three splicing patterns for both of them, it was conformed by sequencing that there were no difference between wild-type and mutant-type. So the mutant-type do not affect splicing.
5. c.1874A>C: The wild type and the mutant type both have one band with the same size. The sequencing results show that both wild type and mutant type were normal spliced bands and the splicing method was “Exon9-Exon10 (148bp)-Exon11”.
6. c.2125-3C>G: The wild type had one band which splicing method was “Exon11-Exon12 (129bp). By contrast, the mutant type had one smaller band which had a 22bp deletion on the left side of Exon12 and the splicing method was “Exon11-△Exon12(107bp).The specific splicing results are shown in Figure 3.

**Figure 2.**
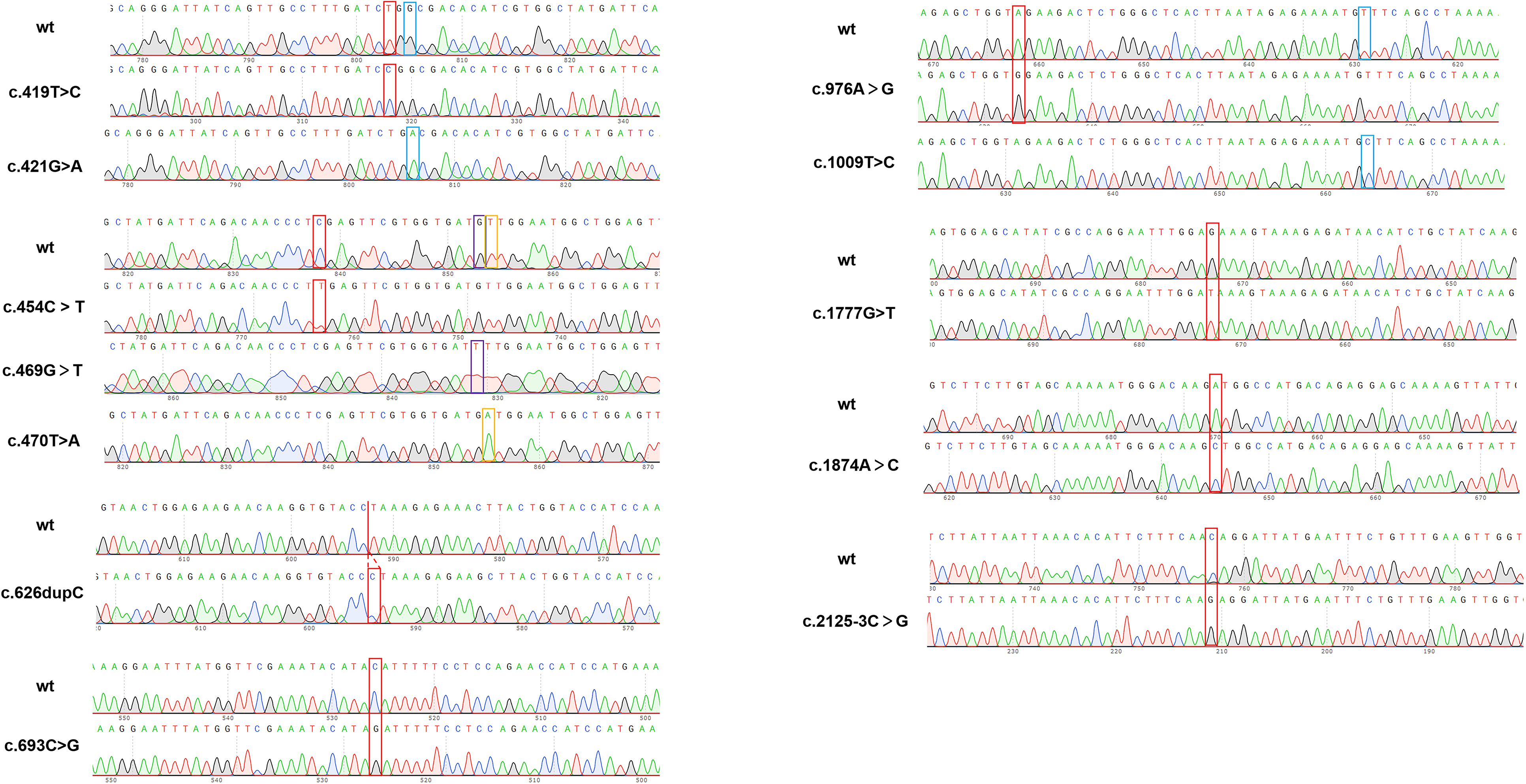
Minigene construction sequencing map: the upper is wild-type, and the bottom is mutant-type.

**Figure 3.**
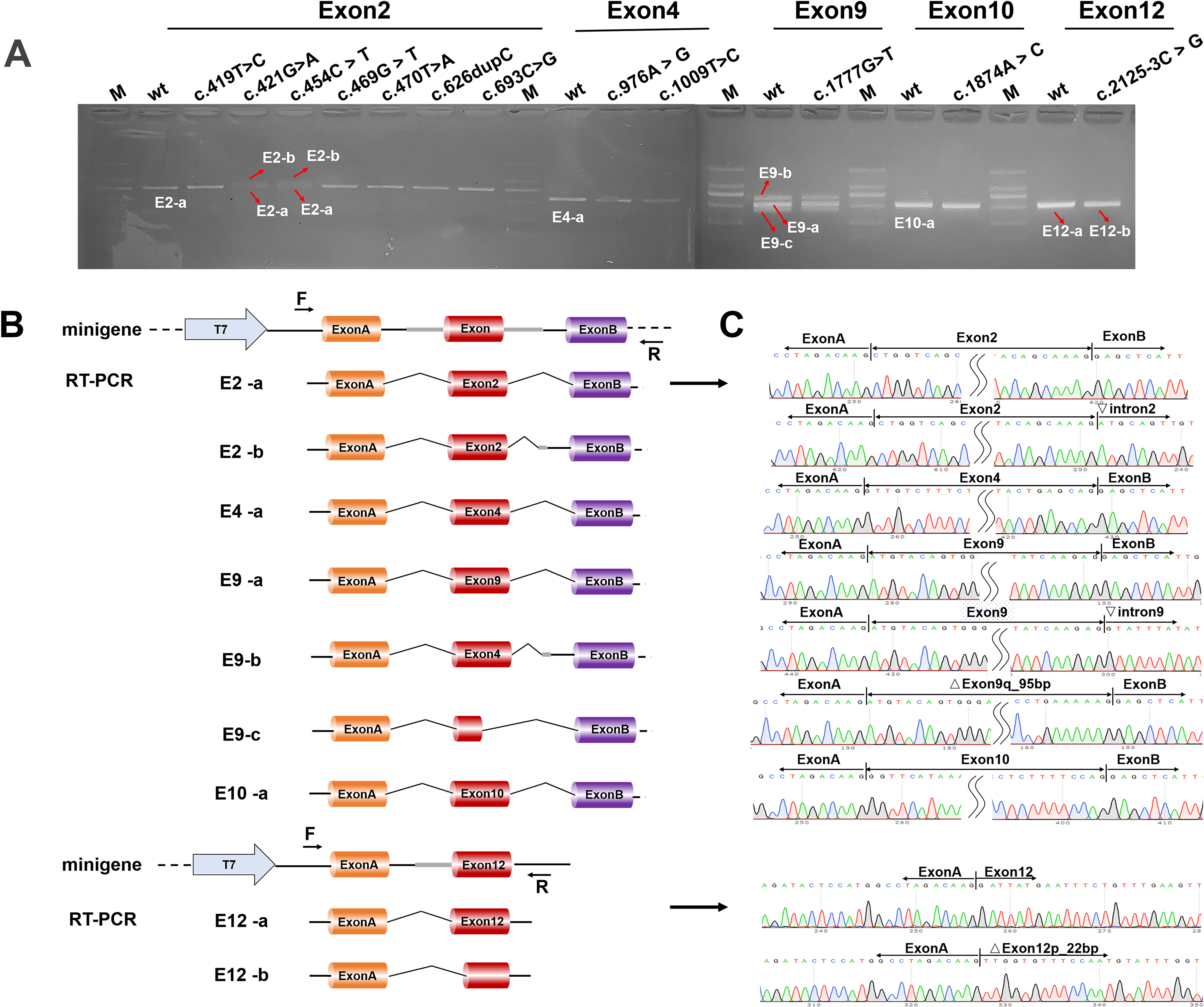
Transcription analysis results. A: RT-PCR showed different bands on the gel which are labeled as a, b, c, respectively; B: Schematic diagram of minigene construction and splicing; C: Sequencing results corresponding to spliced bands. Total 12 mutations were detected. Among them, just three mutations (c.454C>T, c.421G>A and c.2125-3C>G) had been shown to affect mRNA splicing.

## Discussion

With the decrease of high-throughput sequencing cost, the parents of the children with suspected IMD in our Newborn Screening Center are more likely to accept genetic test. However, the increase in the number of cases has also brought about the increase in the number of variant of uncertain significance (VUS) sites, causing a lot of confusion to clinical doctors and patients.

The judgment of pathogenic sites in gene detection reports is based on current literatures, and scientific research is a process with continuous progress. At present, the pathogenicity of mutation sites is generally judged by each testing agency according to the ACMG/AMP guidelines. As for the pathogenicity judgment of mutation sites, more pathogenicity evidences may be found with detailed investigations, so that the initial judgment would be revised. For example, for a VUS related to autosomal recessive hereditary disease, if the subsequent pedigree verification results indicate that it is a de novo mutation (the genetic relationship between the samples is consistent after identification), and the symptoms of the subject are highly consistent with the phenotype of the disease caused by the mutation of the gene, the pathogenicity of the mutation can be upgraded from VUS to a likely pathogenic mutation. Other evidences to support the pathogenicity, including co-separation with phenotype in familial cases, functional verification, update of population database, can be further obtained. The testing agency should establish the regulation to perform regular retrospective analysis of the pathogenicity of the mutation sites. If the level of pathogenicity is adjusted due to new evidences, a correction report should be issued to the subject in time according to the results of re-analysis. In the case reports of isobutyrylglycineuria in recent years, VUS variation is very common. In principle, the literature used to evaluate the pathogenicity of mutation sites in these cases should be evaluated and strictly controlled. It cannot be simply considered that the mutation detected in patients reported in the literature is pathogenic mutation. The research methods, detection scope and whether functional verification is described in the literature should be included in the parameters of pathogenicity evaluation. To judge the pathogenicity of mutation sites, as many data as possible should be referred. In this study, 12 VUS mutations are selected according to the local suspected population. The minigene results confirm that only 3 VUS mutations affect mRNA splicing. Due to the limitations of the study, the other 9 sites can not be classified as “benign” sites. However, based on the results of this study, even if the other 9 sites are pathogenic sites, their pathogenic mechanisms are not due to splicing changes, which are not completely consistent with the software predictions.

Our study on MUT gene could provide an example to demonstrate that the conclusion on suspected pathogenic sites is not completely correct and needs further verification. Especially in the case of primary screening diseases such as methylmalonic acidemia and argininemia, VUS sites will interfere with the prenatal diagnosis and prenatal consultation. Once the VUS sites are not the actual pathogenic sites, it may lead to misdiagnosis of prenatal diagnosis, and then bring pain to the family members of the patients and medical disputes between doctor and patient . Therefore, we want to emphasize again that the literature used to evaluate the pathogenicity of mutation sites should be evaluated and strictly controlled. Combined with the biochemical test and other data, the clinicians should comprehensively interpret the genetic test report to the family members of the children.

## Declarations

### Ethics approval and consent to participate

This project received ethical approval from the Nanjing Maternity and Child Health Care Hospital for data and information. All procedures performed in studies involving human participants were in accord ance with the Nanjing Maternity and Child Health Care Hospital and with the 1964 Helsinki declaration and its later amendments or comparable ethical standards. The parents of the study participants signed a written informed consent form to participate in this study.

### Consent for publication

Parents gave their written consent to participate in this study and to publication, which was approved by the institutional ethics committee of Nanjing Maternity and Child Health Care Hospital.

### Avalilability of data and materials

All data generated or analysed during this study are available from the corresponding author on reasonable request.

### Authors’ contributions

WYY led the review process, detected the sample, analyzed sequencing data, drafted the initial manuscript and extracted data; and SY reviewed all articles; JT is responsible for the overall content. All authors read and approved the final manuscript.

## Acknowledgments

Not applicable

